# TaqMan quantitative real-time PCR for detecting *Avipoxvirus* DNA in various sample types from hummingbirds

**DOI:** 10.1101/2020.03.09.983460

**Authors:** Hanna E. Baek, Ravinder N. Sehgal, Ruta R. Bandivadekar, Pranav Pandit, Michelle Mah, Lisa A. Tell

## Abstract

**Background:** Avian pox is a viral disease documented in a wide range of bird species. Disease related detrimental effects can cause dyspnea and dysphagia, therefore birds with high metabolic requirements, such as hummingbirds, are especially vulnerable. Hummingbirds have a strong presence in California, especially in urban environments; however, little is understood regarding the impact of pox virus on hummingbird populations. Diagnosing pox infections relies on obtaining a tissue biopsy that poses significant bird risks and field challenges. Understanding the ecology of hummingbird pox viral infections could be advanced by a minimally invasive ante-mortem diagnostic method. This study’s goal was to address this gap in understanding if pox infections can be diagnosed using integumentary system samples besides tissue biopsies. To meet this goal, we tested multiple integumentary sample types and tested them using a quantitative real-time PCR assay. A secondary study goal was to determine which sample types (ranging from minimally to highly invasive sampling) were optimal for identifying infected birds.

**Methodology/Principal Findings:** Lesion tissue, pectoral muscle, feathers, toenail, blood, and swabs (both lesion tissue and non-lesion tissues) were taken from live birds and carcasses of two species of hummingbirds found in California. To maximize successful diagnosis, especially for samples with low viral load, a real-time quantitative PCR assay was developed for detecting the hummingbird-specific *Avipoxvirus* 4b core protein gene. *Avipoxvirus* DNA was successfully amplified from all sample types across 27 individuals. Our results were then compared to those of conventional PCR. Comparisons were also made between sample types utilizing lesion tissue samples as the gold standard.

**Conclusions/Significance:** Hummingbird avian pox can be diagnosed without relying on tissue biopsies. Feather samples can be used for diagnosing infected birds and reduces sampling risk. A real-time PCR assay detected viral DNA in various integumentary system sample types and could be used for studying hummingbird disease ecology in the future.

## Introduction

Avian pox is a disease caused by strains of the genus *Avipoxvirus* and can manifest in a cutaneous (dry) and a diphtheritic (wet) form [1]. With the dry form of pox infections, wart-like growths form primarily on non-feathered body regions and are relatively easy to detect. In contrast, the wet form of a pox infection is more difficult to visually detect in a free-ranging bird, as it is characterized by growths on the mucosal membranes in the mouth, esophagus, and lungs [2]. Both forms of pox can cause respiratory or alimentary tract compromise that can ultimately lead to mortality of infected birds [3]. Avian pox can infect a wide range of bird species, including hummingbirds [4]. Hummingbirds appear to be especially vulnerable to the effects of the pox virus because of their high metabolic requirements [5]. Understanding how avian pox impacts hummingbird populations is important since this iconic pollinating avian species can be indicators of environmental health. As such, information about hummingbird diseases can inform us about the general health of the environments they inhabit [5].

Avian pox has traditionally been diagnosed by histological analyses [1,4] and electron microscopy [6] of tissue samples from lesions found on skin surfaces. Though histological analysis is a reliable method for pox diagnosis, it is difficult to do on frozen samples and Bollinger bodies, intracytoplasmic inclusion bodies found in the tissues of pox infected birds [7], are not always found [8]. Additionally, histology requires a tissue biopsy. Taking tissue samples of lesions for analyses is a reliable method for detecting avian pox; however, tissue biopsies taken from live birds present several challenges. Tissue biopsies requires anesthesia and presents the risk of significant hemorrhage or open wounds. Thus, it is important to consider alternative methods for diagnosing avian pox in live birds.

Since tissue biopsies present challenges for avian pox diagnosis in live birds, use of different, less invasive sample types for diagnosis would be beneficial. Since avian pox is a disease that primarily targets the integumentary system [9], we hypothesized that it would be possible to detect avian pox in other sample types that are components of the integumentary system. Taking different sample types would allow for minimal animal harm, especially in the case of field sampling; however, a method for testing the different samples must be developed for reliable diagnosis of avian pox infection.

Polymerase chain reaction (PCR) testing has been used to detect many avian pathogens, such as avian malaria [10,11,12] and avian infectious bronchitis [13]. PCR targets and amplifies specific regions of the pathogen’s genome, which may allow for detection even when the disease is asymptomatic or cannot be diagnosed using histopathology [10] or viral isolation [14]. PCR has been used to diagnose avian pox infections as avian pox is a DNA virus [15]. Our colleagues [2] described a PCR protocol that amplified a 578-bp fragment of fowlpox virus (FPV) from skin tissue samples and respiratory swabs taken from chickens showing signs of pox infection [2]. Our colleagues [16] used a multiplex PCR protocol that could detect both *Avipoxvirus* and papillomavirus infections. Using superficial skin swabs from birds in the field and of preserved museum skin specimens that demonstrated symptoms of viral infection, they reported that detection of multiple strains of both *Avipoxvirus* and papillomavirus is possible through PCR [16]. This protocol was also used by our colleagues [17]; where, in addition to superficial skin swabs, they also tested blood and tissue samples taken from symptomatic wild birds and found that swab and tissues samples generated significantly more avian pox positives than blood samples [17].

In addition to conventional PCR, another method for diagnosing avian pox infections is quantitative real-time PCR. Real-time PCR allows for a quantitative analysis of pathogens without the need for additional diagnostic tests, such as with gel electrophoresis for conventional PCR. As a complementary method to conventional PCR, our colleagues [18] developed a real-time PCR protocol for detecting avian pox in archived blood samples from Hawai’i Amakihi that were confirmed or suspected (wart-like lesions) infections. Cases were confirmed either from the successful culturing of *Avipoxvirus* or through conventional PCR testing. The protocol demonstrated that real-time PCR could be used to positively identify avian pox infections and estimate viral load [18]. As such, real-time PCR may be a useful tool for detecting avian pox infection in hummingbirds as it is a more sensitive assay for detecting viral particles and can be used to detect pox viral DNA in non-lesion tissue samples, which are likely to have lower viral loads.

Although it has been shown that avian pox can be detected via conventional PCR [2,16,17] and real-time PCR [18] in other bird species, a real-time PCR method for detecting the specific strain of *Avipoxvirus* found in hummingbirds has not been developed. Our colleagues [18] screened their samples from Hawai’i honeycreepers using a real-time PCR protocol that amplifies a segment of the *Avipoxvirus* 4b core protein gene. They successfully amplified *Avipoxvirus* DNA in several of their samples, but in their analyses, the Hawaiian strain clusters with canarypox [19], which is distinct from the strain of pox found in hummingbirds [4]. The study conducted by our colleagues [4] used a protocol that amplified a segment of the 4b core protein gene of FPV, which allowed for confirmation of avian pox infection. However, the authors also found that hummingbirds seemed to be infected with a strain not found in other surveyed bird species; this confirms the results in a study [20] that found that poxviruses infecting different species of birds demonstrated considerable variation. Our colleagues [4] found that the pox virus found in hummingbirds seems to cluster with pox viruses isolated from avian species for which a specific diagnostic PCR protocol has not been developed. Thus, in order to most accurately screen hummingbird for pox infection, a protocol that is specific to the strain of avian pox found in hummingbirds would be beneficial.

Developing an accurate method for detecting pox infection without taking tissue biopsies would allow for field studies of *Avipoxvirus* infections. By analyzing and comparing the results from different sample types using real-time PCR as a complement to conventional PCR, it can be determined which sample type might be optimal for screening for pox infections and would allow researchers to prioritize sample collection when in the field or laboratory.

The goals of this study were to 1) determine if pox infection could be diagnosed without a tissue biopsy and 2) determine which integumentary system sample types allow for optimal screening of hummingbirds for pox infection. We describe the development of a quantitative real-time PCR protocol for detecting *Avipoxvirus* in a variety of sample types taken from Anna’s Hummingbirds (*Calypte anna*; ANHU) and a *Selasphorus* Hummingbird (*Selasphorus* spp.; SEHU) to determine if avian pox could be diagnosed without a tissue biopsy. We compared the results of the real-time PCR to those from conventional PCR testing to determine the effectiveness of the real-time PCR protocol. Assessment of the results from different sample types in comparison to lesion tissue samples were made to determine the least invasive sample type that can be taken to assess populations disease prevalence. Additionally, we evaluated the relatedness of the pox viruses found in hummingbirds.

## Materials and Methods

All research within the scope of this study was conducted with permit or committee approval from the United States Fish and Wildlife Service (Permit: MB55944B-2), United States Geological Survey Bird Banding Laboratory (Permit: 23947), California Department of Fish and Wildlife (Permit: SC-013066), and the UC Davis Institutional Animal Care and Use Committee (Protocol: 20355).

### Sample Collection

Various integument samples from 27 individual hummingbirds (ANHU *n=26*; SEHU *n=1*; Table 1), either carcasses or field-caught, were collected. Due to missing feathers, the *Selasphorus* hummingbird could only be identified to genus. Carcasses (*n=19 birds*) were collected from California rehabilitation centers where birds did not survive the rehabilitation process. Some birds (*n=8*) were euthanized during live sampling since they were considered unfit for survival owing to heavy pox infections.

**Table 1.** Summary of sample types (N= number of samples; percentage) taken from Anna’s Hummingbirds (n= 26) and a *Selasphorus* Hummingbird where *Avipoxvirus* was detected by conventional and real-time polymerase chain reaction assay.

| Sample Type | N | Conventional PCR-positive<br>for *Avipoxvirus* | Real-time<br>PCR-positive<br>for *Avipoxvirus* |
| --- | --- | --- | --- |
| **Tissue: Lesions** | 43 | 43 (100%) | 41 (95%) |
| **Tissue: Pectoral**<br>**Muscle** | 26 | 23 (88%) | 26 (100%) |
| **Blood** | 7 | 6 (86%) | 7 (100%) |
| **Toenails** | 29 | 26 (90%) | 27 (93%) |
| **Feathers: Retrices** | 32 | 26 (81%) | 28 (88%) |
| **Feathers: Remiges** | 2 | 2 (100%) | 2 (100%) |
| **Feathers: Contour** | 31 | 24 (77%) | 31 (100%) |
| **Swab (CTA\*):**<br>**Lesion Tissue** | 42 | 42 (100%) | 42 (100%) |
| **Swab (CTA): Non-**<br>**Lesion Tissue** | 15 | 12 (80%) | 12 (80%) |
| **Swab (FTA Card):**<br>**Lesion Tissue** | 1 | 1 (100%) | 1 (100%) |

\*CTA= cotton tipped applicator

To compare samples taken ante- and post-mortem, feather (contour and rectrices), toenail, and lesioned tissue samples were collected while the birds (n=6) were alive and similar samples were collected post-mortem (Supplementary Tables 2 and 3). Blood samples (*n=7 birds*) were taken via clipping the distal 10% of the toenail and collected onto FTA paper [21]. A swab of lesioned tissue was collected from the foot of a live bird using FTA paper [21].

From carcasses, feather, toenail, lesion tissue, and pectoral muscle tissue samples were collected. In addition, swabs of tissues with lesions and no lesions were taken. Although pectoral muscle is not considered to be part of the integumentary system, this sample type was included since it can easily be harvested from specimens destined to be study skins. For toenail samples, approximately 10% of the toenail was sampled from the carcasses. Tissue samples were taken from lesions on the wings, feet, eyes, or bill. For birds with lesions on multiple regions, tissue samples were taken from each region using a sterile no. 15 scalpel blade. Lesion swabs were obtained using sterile cotton-tipped applicator (CTA) swabs soaked in saline solution and pectoral muscle tissue (0.2-0.4 cm diameter) was taken from each carcass using a dermal punch (Miltex Inc., York, Pennsylvania, USA, catalog #s MLTX33-31 and MLTX33-34, respectively). Several different sample types were taken from each individual to increase the likelihood of detecting pox viral DNA as well as to determine which sample types could be reliably used to test for pox infection.

### Samples and DNA Extraction

For the initial TaqMan protocol development, only contour feather and lesion tissue samples were extracted. Approximately 5 mg of lesion tissue or five feathers was added to a 96-well deep-well grinding block (Greiner Bio-One, Monroe, North Carolina, USA, cat #780215) with 600 µl of ATL Buffer, 60 µl of Proteinase K (Qiagen, Valencia, California, USA), and two stainless-steel beads (Fisher Scientific, Waltham, Massachusetts, USA, cat #02-215-512). The grinding block was sealed, then the samples were pulverized in a 2010 Geno/Grinder homogenizer (SPEX SamplePrep, Metuchen, New Jersey, USA) at 1,750 rpm for 2.5 min. Lysate was incubated for 15 min at 56° C. 200 µl of lysate was removed and used for total nucleic acid (TNA) extraction. TNA extraction was performed on a semi-automated extraction system (QIAamp 96 DNA QIAcube HT Kit, QIAcube; Qiagen, Valencia, California, USA) according to manufacturer’s instructions and eluted in 100 µl of diethylpyrocarbonate (DEPC)-treated water. These extraction protocols were modified for sample analysis as detailed later.

A total of 228 samples were taken from various anatomical sites of 27 individual hummingbirds. DNA was extracted from 43 lesion tissue samples where abnormalities were observed on anatomic regions (wings, feet, bills), seven blood samples, 29 toenail samples, 26 pectoral muscle tissue samples, 32 tail feathers (rectrices), 31 contour feather samples, 42 lesion tissue swabs, 15 non-lesion swabs, two wing feathers (remiges), and one FTA swab of lesion tissue (Supplementary Table 2). Some ante-mortem samples were taken to test for the likelihood of viral contamination and were compared to samples taken post-mortem (Supplementary Table 3). For DNA extraction of blood samples, a small piece of Whatman FTA paper (GE Healthcare, Chicago, Illinois, USA) with blood was cut using sterilized dissection scissors (Thermo Fisher Scientific, Carlsbad, California, USA; Stainless steel). The same method was used for DNA extraction of the FTA swab of lesion tissue. For rectrices and remiges, DNA was extracted from feather sheath bases, which were cut using sterilized dissection scissors (Thermo Fisher Scientific, Carlsbad, California, USA; Stainless steel). DNA was also extracted from four whole contour feathers taken from the pectoral region and from toenail clips. DNA was extracted from each sample taken from captured birds and carcasses using the Wizard Genomic DNA Purification kit (Promega Corp., Madison, Wisconsin, USA) according to the manufacturer’s instructions and eluted in 125 µl of DEPC-treated water. All samples taken from carcasses and live birds were stored at -80° C for up to one year before DNA was extracted.

Extracted DNA samples were analyzed using the Qubit 2.0 Fluorometer (Thermo Fisher Scientific, Carlsbad, California, USA) to ensure successful extraction of DNA. These extracted DNA samples were stored at -20° C until PCR analysis.

### Conventional PCR Amplification of Avipoxvirus DNA and Sequencing

Extracted DNA was tested via PCR to amplify a section of the *Avipoxvirus* 4b core protein gene, using a protocol adapted from a previously published study [4]. 4 µl of extracted DNA were used in 25 µl reactions containing 5 µl 5x green PCR buffer, 2.5 µl MgCl_2_ (1.0-4.0mM), 0.5 µl DNTP, 1 µl each of forward and reverse primers [4], 0.125 µl GoTaq Flexi DNA Polymerase (Promega Corp., Madison, Wisconsin, USA), and 10.875 µl Ultrapure water. The forward and reverse primers used in these reactions were those used by our colleagues [4]; however, all other ingredients of the reactions were adjusted for use with GoTaq Flexi DNA Polymerase (Promega Corp., Madison, Wisconsin, USA). Reactions were run through an initial denaturing period of 95° C for 5 min, followed by 40 cycles of 95° C for 30 sec, 50° C for 30 sec, and 72° C for 1 min, and the final step of 72° C for 7 min [4]. Products were visualized on a 1.8% agarose gel. The PCR products of samples that tested positive, one sample per bird (*n=27*), when run through conventional PCR were sent to Elim Biopharmaceuticals, Inc (Hayward, California, USA) for sequencing. The returned nucleotide sequences were manipulated and aligned using Geneious® version 11.1.5 (Biomatters, Inc., San Diego, California, USA), then searched through the National Center for Biotechnology Information Basic Local Alignment Search Tool (BLAST).

### Real-time PCR Assay Development, Validation, and Sample Analysis

The assay was designed to target the 4b core protein gene of *Avipoxvirus* (AAPV). Two primers (vAAPV-124f ACGTCAACTCATGACTGGCAAT and vAAPV-246r TCTCATAACTCGAATAAGATCTTGTATCG) and an internal hydrolysis probe (vAAPV-159p-FAM-AGACGCAGACGCTATA-MGB, 5’ end, reporter dye FAM (6-carboxyfluorescein), 3’ end, quencher dye NFQMGB (Non-Fluorescent Quencher Minor Grove Binding)) were designed using Primer Express Software (Thermo Fisher Scientific, Carlsbad, California, USA). The 123 base pair amplicon was entered into BLAST (NCBI) to confirm unique detection.

In the initial development of the assay, each real-time PCR reaction contained 20X primers and probe with a final concentration of 400nM for each primer and 80nM for the probe, 7 µl of commercially available PCR master mix (TaqMan Universal PCR Master Mix, Thermo Fisher Scientific, Carlsbad, California, USA, cat #4318157) containing 10mL Tris-HCl (pH 8.3), 50mM KCl, 5mM MgCl_2_, 2.5 mM deoxynucleotide triphosphates, 0.625U AmpliTaq Gold DNA polymerase per reaction, 0.25 U AmpErase UNG per reaction, and 5 µl of diluted extracted DNA. The real-time PCR was performed using the ABI PRISM 7900 HTA FAST (Thermo Fisher Scientific, Carlsbad, California, USA). The following amplification conditions were used: 50° C for 2 min, 95° C for 10 min, 40 cycles of 95° C for 15 sec and 60° C for 1 min. Fluorescent signals were collected during the annealing phase and Cq (quantitation cycle) values were extracted with a threshold of 0.2 and baseline values of 3-10. Cq values are the number of cycles required for the fluorescent signal to exceed the background level. Cq values are inverse to the copies of target DNA in a sample (i.e. lower Cq values indicated high amounts of target sequence). A no template control (DEPC-treated water) was run with all assays to ensure absence of non-specific binding of the primers and probes. A reference gene, eukaryotic 18S assay (Hs99999901_s1, Applied Biosystems, Thermo Fisher Scientific, Carlsbad, California, USA), was run with each sample to confirm successful DNA extraction and lack of PCR inhibitors. Positive controls (AAPV plasmid and pooled DNA for 18S) were run with their respective assay to ensure the assay was working properly.

The AAPV assay was validated for efficiency and sensitivity by running a 10-fold standard curve (Figure 1) in triplicate of serial dilutions made from PCR2.1 plasmid DNA (Eurofins Genomics LLC, Louisville, Kentucky, USA) containing the AAPV amplicon. The real-time PCR assay was found to be 94.5% efficient and sensitive enough to detect as few as 10 copies of the target gene per qPCR reaction (R^2^ value = 0.998).

**Figure 1.**
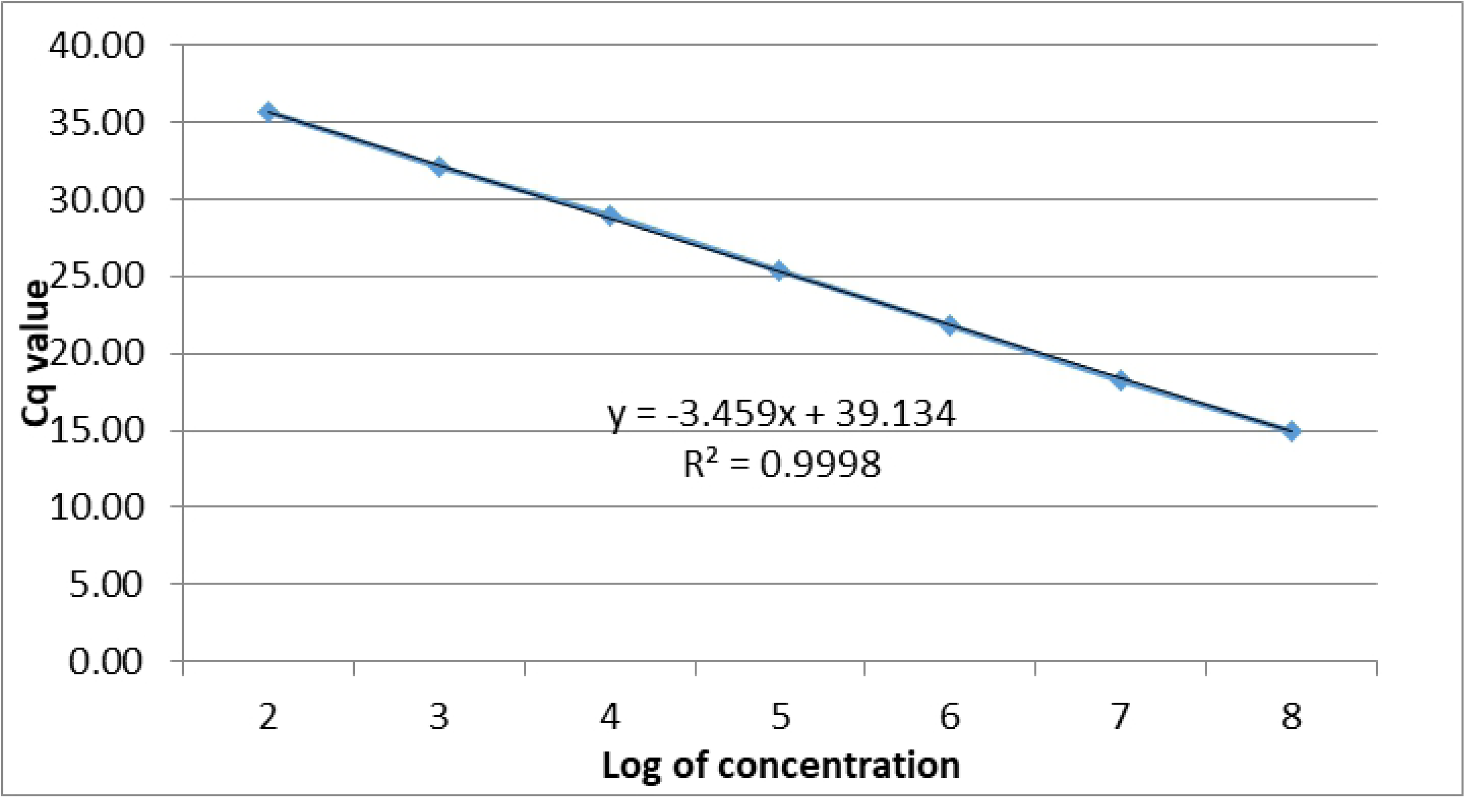
Standard curve developed for absolute quantification of viral DNA copies developed through a triplicate test of 10-fold serial dilutions of *Avipoxvirus* plasmid.

Extracted DNA was tested via real-time PCR to amplify a section of the *Avipoxvirus* 4b core protein gene. Samples were analyzed using the validated assay though the following minor modifications were made. A different commercially available PCR master mix was used (TaqMan Fast Advanced Master Mix, Thermo Fisher Scientific, Carlsbad, California, USA, cat #4444557), but the same concentrations of reagents were used (7 µl master mix: 5 µl extracted DNA). Reactions were run on a different system, CFX 96 Touch Real-Time PCR Detection System (Bio-Rad, Hercules, California, USA), under the same amplification conditions as the validated protocol: 50° C for 2 min, 95° C for 10 min, 40 cycles of 95° C for 15 sec and 60° C for 1 min. A positive control (AAPV plasmid) was run with all assays to ensure the assay was working properly; however, pooled DNA for 18S was not included in the assay during sample analysis. A no template control (purified water) was also run with all assays to ensure the absence of non-specific binding of the primers and probe. Fluorescence signals were collected during amplification and Cq values were extracted for each sample.

### Absolute Quantification of Viral DNA

To quantify the number of copies of AAPV target genes in the samples, a plasmid standard curve was prepared in triplicate using 10-fold serial dilutions. To determine molecules/μl, the following formula was used:

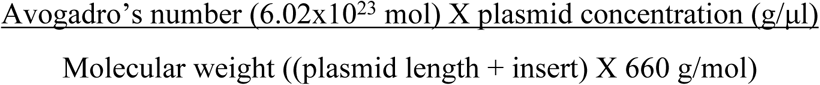

To determine absolute number (*abs*), the following formula was used: Log^10^((Cq-*y*) ÷ *s*), where *y* is the y-intercept and *s* is slope obtained from a plotted standard curve (Figure 1). To determine the number of copies per well, the following formula was used: 10^*abs*^ ÷ 2. The copy number per well (1µl DNA) was divided by two since there are two copies of the gene per cell.

### Assessment of Reliability of Real-time PCR in Detecting Positives

Cohen’s kappa statistics (*k*) was used to evaluate the agreement between conventional and real-time PCR results for each sample type. The closer the value of *k* to 1, the more the agreement between the two tests. To understand the performance of the newly developed real-time PCR, we estimated the sensitivity, positive predictive value (PPV), and F-1 (harmonic mean of sensitivity and PPV) statistic for each sample type [22]. For this analysis, the true status (positive or negative for pox) of an individual bird was determined based on the results of all tests (both conventional and real-time PCR) performed on samples taken from the bird. By this criterion, a bird was assumed to be truly positive if any sample was detected positive for either conventional PCR or real-time PCR. The *k* statistic was also calculated to understand the agreement between different sample types for the real-time PCR.

## Results

Of 228 samples that were fluorometrically analyzed, 174 demonstrated double-stranded DNA concentrations of at least 0.5 ng/mL and the remaining 54 demonstrated concentrations lower than 0.5 ng/mL.

All 228 samples were tested using the conventional PCR assay for the presence of avian pox virus, of which 90% (*n=205*) tested positive (Table 1). Lesion tissue samples (*n=43/43*) as well as lesion tissue swabs (*n=42/42*) were most commonly positive (100%), but all other sample types tested positive as well (Table 1). 100% of remiges (*n=2/2*) and FTA swabs of lesion tissue (*n=1/1*) also tested positive; however, the low sample size must be considered. Toenails and pectoral muscle tissue samples were also positive at a high frequency, with 90% (*n=26/29*) of toenail samples and 88% (*n=23/26*) of pectoral muscle tissue samples determined as positive for pox virus using the conventional PCR assay. Contour feathers were the least likely to be positive at 77% (*n=24/31*) when analyzed using the conventional PCR assay but were very likely (100%) to be positive when tested with the real-time PCR assay (*n=31/31*).

From 27 birds, PCR product sequences from 24 birds were 100% identical to the sequence for pox virus in Anna’s Hummingbirds that was previously published (GenBank accession JX418296) by our colleagues [4]. For the three remaining birds, sequences from two were insufficient to accurately determine nucleotide differences. With the last individual Anna’s Hummingbird, we may have encountered a distinct lineage with three base pairs differing from the common consensus sequence.

All 228 samples were analyzed through the AAPV assay and the average Cq values per sample type were calculated (Table 2). The real-time PCR found that 95% (*n=217*) of 228 samples tested positive for *Avipoxvirus.* Of the samples that were negative for *Avipoxvirus* (*n=11*), five contained low DNA concentrations (<0.5 ng/mL), which may have resulted in false negatives. To indicate successful viral amplification, we increased the threshold of Cq values used by our colleagues [18] up to 40; however, we classified samples with Cq values of 35 to 40 as having a low viral load. Lesion tissue samples seem to contain the highest viral load as they have the lowest average Cq value; however, lesion tissue swabs, remiges, and rectrices also showed significantly low Cq values as well (Table 2). In general, blood samples showed the highest Cq values (Table 2) but amplification of pox virus DNA was successful in all samples (*n=7*; Table 1).

**Table 2.** Average ± Standard Deviation (range) of quantification cycle (Cq) values by sample type for samples taken from Anna’s Hummingbirds (n=26) and a *Selasphorus* spp. Hummingbird and tested via a real-time polymerase chain reaction assay.

| Sample Type | Average Cq Value |
| --- | --- |
| Tissue: Lesions | $19.50 \pm 4.68$ (14.50-31.12) |
| Tissue: Pectoral Muscle | $31.92 \pm 3.79$ (21.60-38.40) |
| **Blood** | 33.14 ± 3.38 (29.26-38.64) |
| **Toenails** | 29.77 ± 3.83 (20.70-38.63) |
| **Feathers: Rectrices** | 28.85 ± 3.24 (23.59-37.44) |
| **Feathers: Remiges** | 27.22 ± 2.80 (25.24-29.20) |
| **Feathers: Contour** | 27.37 ± 4.06 (23.14-37.79) |
| **Swab (CTA\*): Lesion Tissue** | 21.32 ± 3.52 (15.72-29.63) |
| **Swab (CTA): Non-Lesion Tissue** | 29.17 ± 4.41 (18.49-36.08) |
| **Swab (FTA Card): Lesion Tissue** | 22.26 ± 0 (22.26-22.26) |

\*CTA=cotton tipped applicator

At the individual bird level, both conventional and real-time PCR assays were able to detect pox virus in at least one sample type for all birds. When explored for various sample types, conventional and real-time PCR assays showed high agreement of 89.72% with kappa (*k*) of 0.77 (n = 632). Agreement between conventional and real-time PCR assays for various sample types and their corresponding kappa (*k*) values are shown in Table 3. Real-time and conventional PCR results for lesion swab samples showed perfect agreement in correctly identifying positive samples (n = 42, no *k* was calculated due to lack of observations in d* category, Table 3). Feathers as sample types showed moderate concordance when tested with conventional and real-time PCR assays (*k*=0.449, *n=65 samples*).When anatomic location of samples were analyzed, tail feathers showed the highest concordance with *k* of 0.76 (n =32 samples).

**Table 3.** Assessment of the performance of a real-time PCR assay for detecting *Avipoxvirus* in samples from Anna’s (n=26) and *Selasphorus* spp. (n=1) Hummingbirds. The left part of the table shows confusion matrices, Cohen’s kappa for agreement between the real-time PCR and conventional PCR assays. The right side of the table shows the performance of the real-time PCR assay.

| *Sample Type* | Agreement with conventional PCR assay |  |  |  |  | Real-time PCR assay performance (comparison with true status) |  |  |  |
| --- | --- | --- | --- | --- | --- | --- | --- | --- | --- |
|  | a* | b* | c* | d* | k | Sensitivity | PPV | F-1* | n (number of samples) |
| **Blood** | 6 | 1 | 0 | 0 | 0 | 1 | 1 | 1 | 7 |
| **Feathers** | 53 | 8 | 0 | 4 | 0.449 | 0.94 | 1 | 0.97 | 65 |
| **Toenail** | 25 | 2 | 1 | 1 | 0.345 | 0.93 | 1 | 0.96 | 29 |
| **Lesion FTA Swab** | 1 | 0 | 0 | 0 | N/A | 1 | 1 | 1 | 1 |
| **Lesion Tissue** | 41 | 0 | 2 | 0 | 0 | 0.95 | 1 | 0.98 | 43 |
| **Muscle** | 23 | 3 | 0 | 0 | 0 | 1 | 1 | 1 | 26 |
| **Lesion Tissue Swab** | 42 | 0 | 0 | 0 | 0 | 1 | 1 | 1 | 42 |
| **Non-lesion Swab** | 12 | 0 | 0 | 3 | 1 | 0.8 | 1 | 0.89 | 15 |
| ***Sample types with anatomic location*** |  |  |  |  |  |  |  |  |  |
| ***Feathers*** |  |  |  |  |  |  |  |  |  |
| **Rectrice** | 26 | 2 | 1 | 4 | 0.76 | 0.88 | 1 | 0.93 | 32 |
| **Contour** | 25 | 6 | 0 | 0 | 0 | 1 | 1 | 1 | 31 |
| **Remige** | 2 | 0 | 0 | 0 | N/A | 1 | 1 | 1 | 2 |
| ***Toenail*** |  |  |  |  |  |  |  |  |  |
| **Toenail** | 25 | 2 | 1 | 1 | 0.345 | 0.93 | 1 | 0.96 | 29 |
| ***Tissue*** |  |  |  |  |  |  |  |  |  |
| **Muscle Pectoral** | 23 | 3 | 0 | 0 | 0 | 1 | 1 | 1 | 26 |
| ***Blood*** |  |  |  |  |  |  |  |  |  |
| **Blood** | 6 | 1 | 0 | 0 | 0 | 1 | 1 | 1 | 7 |
| ***Lesion Tissues*** |  |  |  |  |  |  |  |  |  |
| **Lesion Tissue Foot** | 18 | 0 | 0 | 0 | N/A | 1 | 1 | 1 | 18 |
| **Lesion Tissue Beak** | 13 | 0 | 1 | 1 |  | 0.93 | 1 | 0.96 | 14 |
| **Lesion Tissue Wing** | 5 | 0 | 1 | 0 | 0 | 0.83 | 1 | 0.91 | 6 |
| **Lesion Tissue Eye** | 3 | 0 | 0 | 0 | N/A | 1 | 1 | 1 | 3 |
| **Lesion Tissue Keel** | 2 | 0 | 0 | 0 | N/A | 1 | 1 | 1 | 2 |
| ***Lesion Swabs*** |  |  |  |  |  |  |  |  |  |
| **Lesion Swab Wing** | 7 | 0 | 0 | 0 | N/A | 1 | 1 | 1 | 7 |
| **Lesion Swab Beak** | 16 | 0 | 0 | 0 | N/A | 1 | 1 | 1 | 16 |
| **Lesion Swab Foot** | 15 | 0 | 0 | 0 | N/A | 1 | 1 | 1 | 15 |
| **Lesion Swab Keel** | 1 | 0 | 0 | 0 | N/A | 1 | 1 | 1 | 1 |
| **Lesion Swab Eye** | 3 | 0 | 0 | 0 | N/A | 1 | 1 | 1 | 3 |
| ***Non-lesion swabs*** |  |  |  |  |  |  |  |  |  |
| **Non-Lesion Swab Beak** | 5 | 0 | 0 | 1 | 1 | 0.83 | 1 | 0.91 | 6 |
| **Non-Lesion Swab Foot** | 5 | 0 | 0 | 0 | N/A | 1 | 1 | 1 | 5 |
| **Non-Lesion Swab Eye** | 2 | 0 | 0 | 1 | 1 | 0.67 | 1 | 0.8 | 3 |
| **Non-Lesion Swab Wing** | 1 | 0 | 0 | 0 | N/A | 0 | 0 | 0 | 1 |
| ***Other*** |  |  |  |  |  |  |  |  |  |
| **Lesion FTA Swab Foot** | 1 | 0 | 0 | 0 | N/A | 1 | 1 | 1 | 1 |
| ***Lesion vs Swab*** |  |  |  |  |  |  |  |  |  |
| **Lesion Tissue** | 41 | 0 | 2 | 0 | 0 | 0.95 | 1 | 0.98 | 43 |
| **Lesion Swab** | 42 | 0 | 0 | 0 | N/A | 1 | 1 | 1 | 42 |
| **Non-Lesion Swab** | 12 | 0 | 0 | 3 | 1 | 0.8 | 1 | 0.89 | 15 |

\*a: real-time PCR = positive, conventional PCR = positive, b: real-time PCR = negative, conventional PCR = negative, c: real-time PCR = positive, conventional PCR = negative, d: real-time PCR = negative, conventional PCR = positive, F-1 = harmonic mean of precision and recall.

The real-time PCR results showed a high positive predictive value of 1 for all sample types (except non-lesion wing), while the sensitivity (except for non-lesion swabs) ranged from to 1.0. The values varied within sample type when anatomic region was included in the analysis (Table 3). Foot lesion tissue, feather, and pectoral muscle tissue samples showed high sensitivity while the sensitivity of real-time PCR was lower for beak lesion and wing lesion tissue samples (Table 3).

## Discussion

Our results indicate that avian pox can be diagnosed without relying solely on lesion tissue biopsies. Using a variety of integumentary system sample types, pox infections were diagnosed when both conventional and real-time PCR was used. To our knowledge, this is the first time that feather samples have been used to diagnose avian pox infection in any bird species. Likewise, we successfully amplified *Avipoxvirus* DNA from swabs taken from tissue lesions. Our results suggest that pox can be diagnosed with minimally invasive sampling, reducing the risk of stress for birds being handled [23].

In this study, we collected and tested multiple sample types to determine which were optimal for diagnosing pox infection. Lesion tissue was tested as it is the most likely to have high viral load and could be used as a proxy for a “gold standard” for pox detection. Lesion swabs and non-lesion swabs were sampled to determine if virus could be detected on integument surface areas. Feathers and toenail samples were chosen because pox virus targets the integumentary system [9,24,25]. Contour feathers were collected as they are easy to sample and minimally impact the bird if sampling is conservative. Contour feathers were also chosen as a sample type to compare to remiges and rectrices, which are not optimal for sampling, especially during the breeding season given the importance of rectrices during courtship [26]. Toenail samples were tested to optimize ante-mortem diagnosis since toenail clipping is a method for sampling avian blood in the field. Whole blood was taken because this is a sample type that can be taken from live birds. Previous studies have found that blood samples may not be reliable for diagnosing avian pox; however, this could be due to a low concentrations of circulating virus at the time of sampling from a suspected bird [18]. Pectoral muscle tissue was chosen as a sample type, as it can easily be sampled from a carcass intended for a museum collection study skin with minimal damage to the specimen. All tissue (lesion and pectoral muscle), lesion swab, toenail, blood and feather samples showed reasonably high sensitivity and could be considered when testing birds for pox infections.

We tested all samples using both conventional and real-time PCR to determine if there were differences in sensitivity of detecting *Avipoxvirus* DNA. Some samples that were determined negative using conventional PCR were positive when analyzed using real-time PCR testing and returned relatively low Cq values. This finding suggests that real-time PCR has the advantage over conventional PCR in its ability to detect *Avipoxvirus* DNA in samples with low viral load. Specifically, for contour feather samples, real-time PCR yielded a positive result for all birds whereas conventional PCR only detected 77% of the cases. This difference in ability to detect small viral concentrations in samples should also be taken in account when interpreting the predictive values and sensitivity.

Conventional PCR has been used to detect avian pox in the past [2,4] as well as to determine prevalence and genetic diversity of avian pox [27,28]; however, there are still limitations. In analyses using solely conventional PCR, parasite or viral load is difficult to determine [29] and as it relies on downstream visualization, products are handled more than once thus increasing the risk of viral contamination [30]. In addition, compared to real-time PCR, conventional PCR might not be as sensitive for detecting low viral loads. Real-time PCR can be used to determine relative viral load. However, there are also limitations as in some cases, real-time PCR may not detect *Avipoxvirus* DNA in samples taken from pox-positive birds [18]. This was evident in our study, as there were some instances where real-time PCR did not detect avian pox in samples, even those that were found positive using conventional PCR. Despite the limitations, the real-time PCR assay we developed was able to detect very low viral loads (Cq values above 35). Based on our results, real-time PCR holds promise for identifying hummingbirds with pox viral infections using samples, such as feathers, that are taken less invasively. These findings also suggest that asymptomatic birds can be screened for pox by using these sample types.

For this study, all sampled hummingbirds had suspected pox lesions on one or more integumentary regions. Since many samples came from carcasses, there is the possibility of contamination across samples taken from the same individual. We attempted to address this problem by taking ante-mortem samples to compare to post-mortem samples from the same individual (Supplementary Table 3). Our results showed that taking samples ante-mortem had just as much success in diagnosing an infected bird as those taken post-mortem thus making sample contamination post-mortem less likely.

Although hummingbirds are organisms of public interest, there is little known about how avian pox affects hummingbird populations. Hummingbirds are found throughout most of California, the Anna’s Hummingbird particularly, is found in most regions year-round. They can be found in both remote and human-populated areas. With this research, we have expanded our knowledge on how to optimally sample from hummingbirds in order to screen for pox infection without relying on tissue biopsies. With the developed assay, we can begin to answer questions regarding *Avipoxvirus* prevalence in wild hummingbird populations, including how human activities may impact the transmission of this disease. As this assay was developed specifically for hummingbirds, it could also provide a means to help determine the transmission routes of the virus: for example, if the virus is spread by insect vectors or through direct contact transmission.

## Acknowledgements

The authors thank Mrs. Cara Wademan and Mrs. Samantha Barnum for their assistance with developing the real-time PCR protocol. This study was supported in part by generous donations from the Hunter-Jelks Foundation, the Daniel and Susan Gottlieb Foundation, Mr. and Mrs. Thomas Jefferson and Dr. Grant Patrick.

## Supporting Information Captions

**Table 1. Summary of conventional and real-time PCR testing results for *Avipoxvirus* for all sample types taken from individual hummingbirds (n=26 Anna’s Hummingbird and n=1 *Selasphorus* spp. hummingbird).**

**Table 2. Summary of ante-mortem and post-mortem samples collected for *Avipoxvirus* testing from Anna’s Hummingbirds (n=26) and a *Selasphorus* spp. Hummingbird.**

**Table 3. Comparison of results from conventional and real-time polymerase chain reaction testing for birds where the same sample types were taken ante-mortem and post-mortem.**

